# SMU_1361c regulates the oxidative stress response of *Streptococcus mutans*

**DOI:** 10.1101/2023.07.10.548324

**Authors:** Shuxing Yu, Jun Huang, Yaqi Liu, Jing Li, Qiong Zhang, Yan Wang, Tao Gong, Qizhao Ma, Jing Zou, Yuqing Li

## Abstract

Dental caries is the most common chronic infectious diseases around the world and disproportionately affects the marginalized socioeconomic group. *Streptococcus mutans*, considered a primary etiological agent of caries, must depend on the coordinated physiological response to tolerate the oxidative stress generated by commensal species within dental plaque, which is a critical aspect of its pathogenicity. Here, we identified and characterized a novel TetR family regulator SMU_1361c, encoded by the TnSmu2 operon, which appears to be acquired by the bacteria via horizontal genes transfer. Surprisingly, *smu_1361c* functions as a transcriptional repressor to regulate gene expression outside its operon, involved in the oxidative stress response of *S. mutans*. The *smu_1361c* overexpression strain UA159/pDL278-*1361c* was more susceptible to oxidative stress and less competitive against hydrogen peroxide generated by commensal species *Streptococcus gordonii* and *Streptococcus sanguinis*. Transcriptomics analysis revealed that *smu_1361c* overexpression resulted in the significant downregulation of 22 genes mainly belonging to three gene clusters responsible for the oxidative stress response. The conversed DNA binding motif of SMU_1361c was determined by electrophoretic mobility shift and DNase I footprinting assay with purified SMU_1361c protein, therefore, *smu_1361c* is directly involved in gene transcription related to the oxidative stress response. Crucially, our finding provides a new understanding of how *S. mutans* deals with the oxidative stress that is required for pathogenesis and will facilitate the development of new and improved therapeutic approaches for dental caries.

## Author summary

*Streptococcus mutans* is the major organism associated with the development of dental caries, which globally is the most common chronic disease. To persist and survive in biofilms, *S. mutans* must compete with commensal species which occupy the same ecological niche. Here, we uncover a novel molecular mechanism of how TetR family regulator *smu_1361c* encoded by the TnSmu2 genomic island involves in the oxidative stress response through transcriptomics analysis, electrophoretic mobility shift assay, and DNase I footprinting assay. Furthermore, we demonstrated that *smu_1361c* mediates *S. mutans* sensitivity to oxidative stress and competitiveness with commensal streptococci. Therefore, this study has revealed a previously unknown regulation between *smu_1361c* and genes outer its operon and demonstrated the importance of smu_*1361c* in the oxidative stress response and the fitness of S. mutans within the plaque biofilms, which can be exploited as a new therapy to modulate the ecological homeostasis and prevent dental caries.

## Introduction

Dental caries, characterized by the demineralization and subsequent destruction of teeth, is the most prevalent biofilm-associated oral infectious disease globally [1,2]. The pathogenesis of dental caries is considered to be a microbial shift in the biofilm composition; however, *Streptococcus mutans* is the most closely and constantly linked with caries formation [3,4]. Occasionally, *S. mutans* can enter the bloodstream and cause systemic infections, such as infective endocarditis [5–7].

*S. mutans* resides in biofilms on the tooth surface, which is a diverse, multi-species, and dynamic environment that is constantly exposed to severe limitations in nutrition, fluctuations in temperature and pH, and challenges in osmotic and oxidative tensions [8,9]. To survive adverse environmental conditions, bacteria evolve through a wide range of rapid and adaptive responses, including carbohydrate utilization, acid production, and low pH adaption, which are well-studied [10,11]. However, limited information is available regarding oxidative stress response and its impact on the adaptation of *S. mutans*. In a previous transcriptomic study that measured gene transcripts expressed in natural oral biofilm communities, the majority of stress response transcripts (50%–75% of total) were associated with oxidative stress, suggesting that oxidative stress is the leading stressor in the biofilm microbial communities [12,13].

Multiple sources contribute to oxidative stress, including the poisonous effects of reactive oxygen species (ROS) and unfavorable cellular redox potential [14,15]. Damaging ROS, including superoxide anion radical (O2·^−^), hydrogen peroxide (H_2_O_2_), and hydroxyl radical (HO·), are produced inside the bacterial cells when growing in an aerobic environment. O2·^−^ and HO· derives from Fenton chemistry as part of the bacterial oxidative stress pathway [16,17]. ROS are toxic and can react more voluntarily with biological macromolecules, such as cleaving RNA/DNA, and oxidizing proteins and lipids [18,19]. Therefore, the ability of *S. mutans* to cope with oxidative stress is important for their survival and perseverance in dental biofilm microbial communities.

In RNA sequencing (RNA-Seq) analysis, evidence for oxidative stress tolerance is revealed in a large genomic island termed TnSmu2 [20]. Many genes encoded within the TnSmu2 regions are affected in the deletion of *fstH*, *cidB,* or *covR*, which are involved in oxidative stress resistance [21–23]. TnSmu2 is a mobile genetic element commonly acquired through horizontal gene transfer events, which is a selective competitive growth advantage seen under changing environmental conditions that affects the metabolism or pathogenic potential of the organisms [24–26]. The region contains two tetracycline repressor (TetR) family transcriptional regulators, *smu_1349* and *smu_1361c* [27,28]. *smu_1349* is involved in oxidative stress response by regulating the nonribosomal peptide synthetase-polyketide synthase gene cluster [28]. However, the regulatory effect of *smu_1361c* on the oxidative stress tolerance of *S. mutans* is still uncharacterized. Therefore, we suggest a hypothesis that *smu_1361c*, carried on the TnSmu2 genomic island, provides an additional layer of the regulatory mechanism of oxidative stress for *S. mutans*.

Herein, we investigated the function of the transcription factor SMU_1361c, and uncovered its regulatory mechanism on the oxidative stress response. SMU_1361c inhibits the gene expressions of *smu_137-141* by directly binding to their promoters, attributing to the observed defects of *S. mutans* in tolerance against oxidative stress. This study highlights a new link between SMU_1361c from the TnSmu2 regions and the oxidative stress response and provides a new possible goal for the ecological control of dental caries.

## Results

### SMU_1361c increased susceptibility to the oxidative stress response of *S. mutans*

The overexpression and in-frame deletion mutants for *smu_1361c* were constructed to evaluate the function of SMU_1361c. First, we measured the mRNA levels of *smu_1361c* in UA159, UA159/pDL278, UA159/pDL278-*1361c*, and UA159 Δ*1361c*, using quantitative reverse transcription polymerase chain reaction (qRT-PCR). As shown in S1 Fig, the expression level of *1361c* in the strain UA159/pDL278-*1361c* increased significantly, about 50-fold, compared with that of UA159/pDL278, and no *smu_1361c* transcripts were detected in the strain UA159 Δ*1361c*, as expected. These results indicate that both the in-frame deletion mutants for *smu_1361c* were successfully constructed.

We first measured the growth rates of UA159, UA159/pDL278, UA159/pDL278-*1361c*, and UA159 Δ*1361c* under aerobic conditions. No significant differences were observed between these bacterial growths both in the logarithmic growth and stationary phase (data not shown). Furthermore, the effect of *smu_1361c* on other key virulence properties and bacterial fitness was detected. To compare their ability to grow and survive when exposed to hydrogen peroxide, a hydrogen peroxide sensitivity assay was performed. A dose-dependent inhibition was observed when filter disks infused with different hydrogen peroxide concentrations ranging from 20 to 60 mM were placed on a soft agar plate containing UA159 and its derivatives, particularly in the overexpression strain UA159/pDL278-*1361c* (Fig 1). Together, these results suggest that *smu_1361c* plays an important role in oxidative stress response in *S. mutans*.

**Fig 1.**
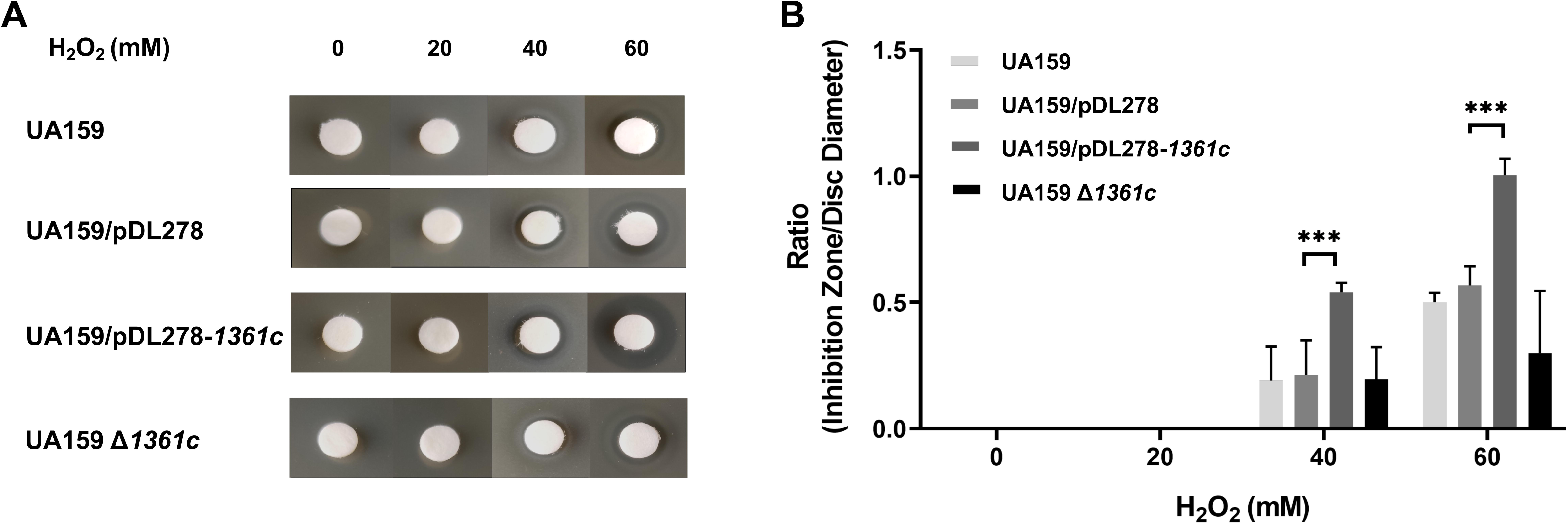
Effect of *smu_1361c* on the oxidative tolerance of *S. mutans*. The plates were overlaid with soft agar containing UA159, UA159/pDL278, UA159/pDL278-*1361c*, and UA159 Δ*1361c* and challenged with filter discs soaked with 0-, 20-, 40-, and 60-mM hydrogen peroxide. The representative plates are shown to demonstrate the differences in inhibition zones (A), which are measured at three different positions and averaged to serve as a single datapoint, and the ratio of inhibition zone to disc diameter was calculated (B). Each experiment was repeated at least thrice. Results are presented as mean ± SD (* *P* < 0.05 or *** *P* < 0.001).

### SMU_1361c impaired bacterial competitiveness of *S. mutans* in the three-species biofilms

The tooth surfaces are colonized by a great number of bacterial species, which normally maintain ecological homeostasis amid different species. Hydrogen peroxide-producing commensal species, including *Streptococcus gordonii* and *Streptococcus sanguinis*, are among many of the early colonizers on tooth surfaces [29,30]. Therefore, the ability of *S. mutans* to contend with oxidative stress generated by hydrogen peroxide is important for its competitiveness with commensal species that occupy the same ecological niche [31–33]. Subsequently, we probed the potential function of *smu_1361c* in regulating the competitiveness of *S. mutans* in oral biofilm communities by creating a three-species biofilms model, comprising *S. mutans*, *S. sanguinis*, and *S. gordonii*. Fluorescent in situ hybridization (FISH) was used to visualize the spatiotemporal interactions within the three-species biofilm communities (Fig 2A). Quantitative analysis of FISH-labeled biofilms showed that the overexpression strain UA159/pDL278-*1361c* was less competitive than the UA159/pDL278 strain against *S. sanguinis* and *S. gordonii* (Fig 2B). No significant competitiveness differences were observed between UA159 and UA159 Δ*1361c* (Fig 2B).

**Fig 2.**
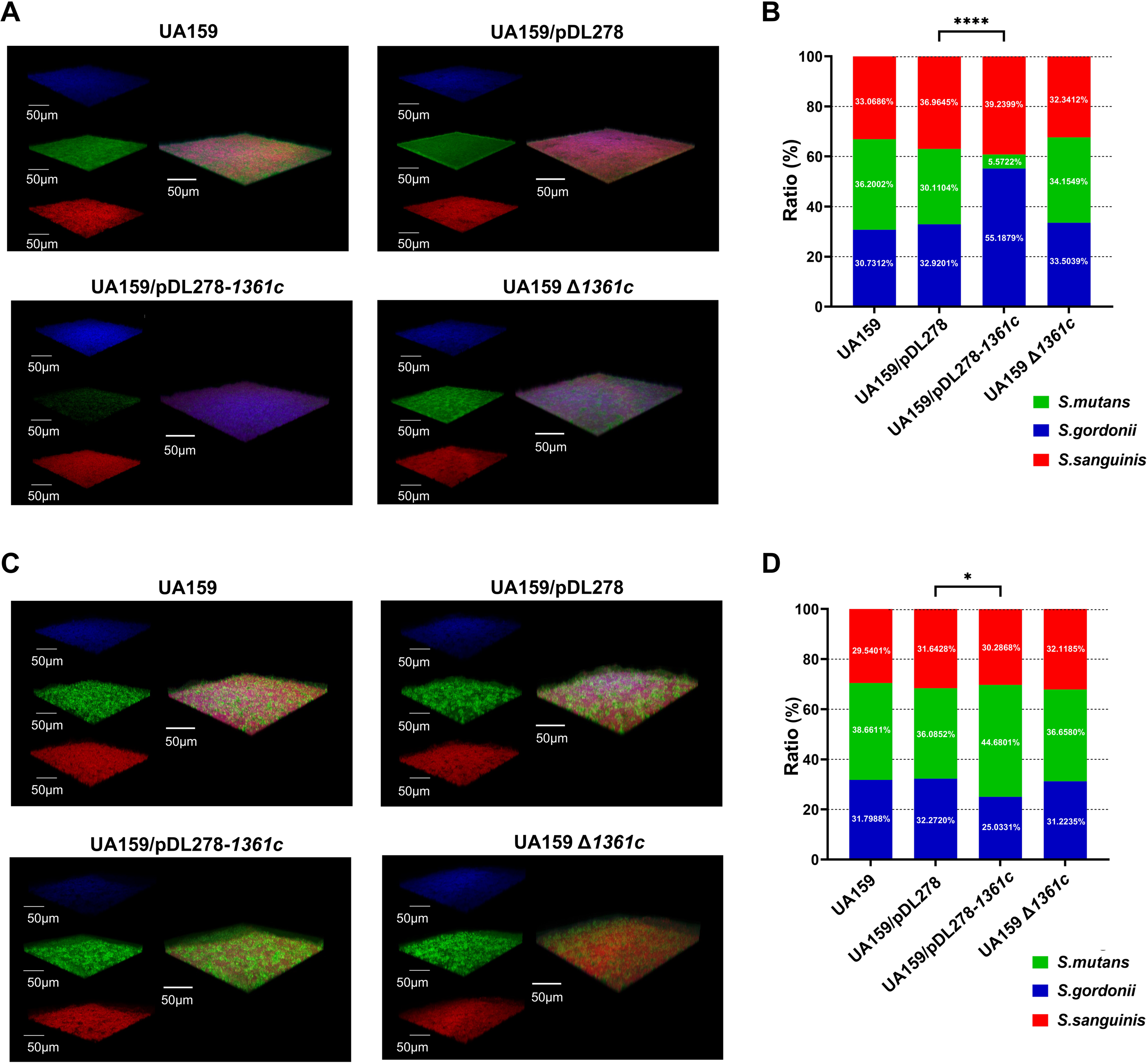
Effect of *smu_1361c* on interspecies competition within the three-species biofilms. *S. mutans* and its derivatives, *S. gordonii*, and *S. sanguinis* were simultaneously cultured in fresh BHIS (A, B) or supplemented with 3 mg/mL catalase (C, D) under aerobic conditions for 24 h and labeled by species-specific fluorescent *in situ* hybridization (FISH) probes. The three-dimensional visualization of three-species biofilms with *S. mutans* (green), *S. gordonii* (blue), and *S. sanguinis* (red) was captured using the confocal laser scanning microscopy (CLSM) at 60× magnification and reconstructed using the IMARIS 7.0.0 (A, C). The ratio of *S. mutans*/*S. gordonii/S. sanguinis* was quantified by the coverage area of each species with Image Pro Plus 6.0 (B, D). The representative images are shown from at least five randomly selected positions of each sample. Results are presented as mean ± SD (* *P* < 0.05 or **** *P* < 0.0001).

To determine whether the less competitiveness of UA159/pDL278-*1361c* is caused by hydrogen peroxide produced by *S. sanguinis* and *S. gordonii*, we detected the effect of hydrogen peroxide on bacterial interactions by adding catalase to the three-species biofilms. As shown in Fig 2C and 2D, there was a superior competitive advantage of UA159/pDL278-*1361c* against *S. sanguinis* and *S. gordonii* over UA159/pDL278; no significant competitive differences were observed between UA159 and UA159 Δ*1361c* against *S. sanguinis* and *S. gordonii*, within three-species biofilms (Fig 2C and 2D). These results indicated that *smu_1361c* impairs the competitiveness of *S. mutans* in biofilm communities owing to the sensitivity to hydrogen peroxide produced by commensal *S. gordonii* and *S. sanguinis*.

### Transcriptomics analysis of the UA159/pDL278-*1361c* strain

To understand the role *smu_1361c* plays globally in regulating oxidative stress response of *S. mutans*, transcriptomics analysis of the UA159/pDL278 and UA159/pDL278-*1361c* strains was performed. In total, 22 significantly downregulated genes (>2-fold) and 14 significantly upregulated genes (>2-fold) were identified in the UA159/pDL278-*1361c* strain compared with UA159 (Fig 3A). Among the up-regulated genes in this range was the gene *smu_1361c*, which further supported the previous qRT-PCR results. Based on the *S. mutans* UA159 genome annotation from the National Center for Biotechnology Information, most of the significantly downregulated genes were mainly associated with energy production and conversion and carbohydrate transport and metabolism, except for some unannotated genes (Fig 3A). The volcano plot illustrates considerably downregulated gene clusters, including *smu_137*-*smu_141*, *smu_764*-*smu_765*, and *smu_127*-*smu_130*, and the highest upregulated gene *smu_1361c* in the UA159/pDL278-*1361c* strain compared with UA159/pDL278 strain (Fig 3B).

**Fig 3.**
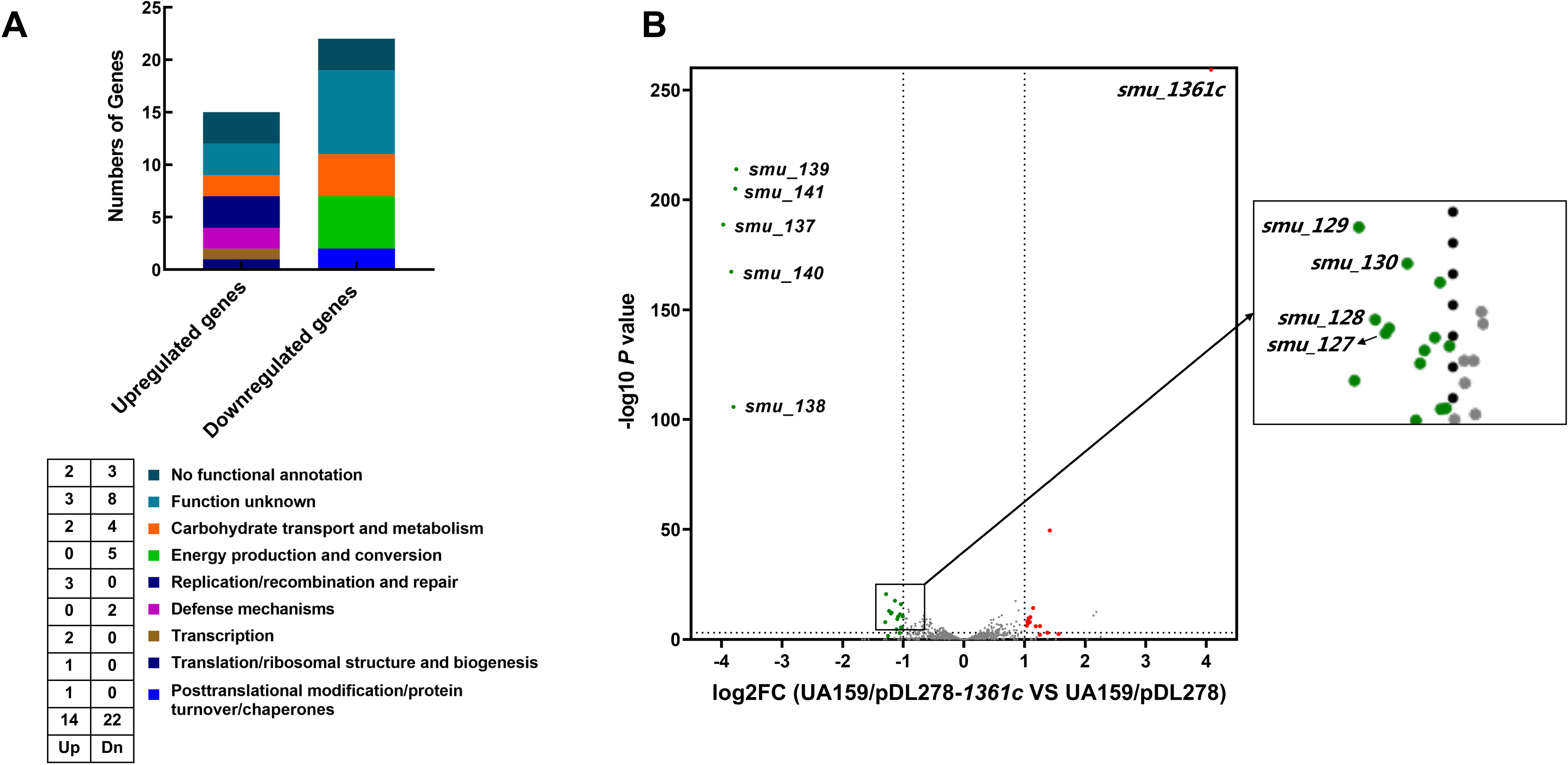
Transcriptomics analysis of UA159/pDL278 and UA159/pDL278-*1361c* strains. (A) Functional classification of differentially expressed genes (DEGs) was performed based on the clusters of orthologous groups (COG) type. (B) The volcano plot illustrates DEGs between UA159/pDL278 and UA159/pDL278-*1361c*.

Among these DEGs, *mleS* (*smu_137*) for a malolactic enzyme, *mleP* (*smu_138*) for malate permease, *gshR* (*smu_140*) for a glutathione reductase known to associate with the oxidative stress response of *S. mutans* were significantly downregulated in the UA159/pDL278-*1361c* strain compared with those in UA159/pDL278. In addition, the gene cluster *smu_127*-*smu_130* was significantly downregulated in UA159/pDL278-*1361c*, which was also recently reported to be involved in the oxidative stress response of *S. mutans* [34]. Significantly, these results indicate that *smu_1361c* may regulate their gene expression by affecting the promoter of these gene clusters (Fig 3).

### SMU_1361c directly bound to the promoters of *smu_137*-*smu_141*

Based on the phenotype results that the UA159/pDL278-*1361c* strain was more sensitive to oxidative stress than the UA159/pDL278 strain, the promoters of *smu_137*-*smu_141* was chosen to study the DNA binding activity and specificity of SMU_1361c and was renamed *smu_137* promoter in subsequent studies, respectively. The *smu_137* promoter (116 bp) was amplified to investigate their interactions with purified recombinant His-SMU_1361c protein using EMSA, and the *smu_136c* ORF fragment (116 bp) was used to exclude the nonspecific bindings of SMU_1361c to DNA substrates (Fig 4A). The mobility shifts were only observed when SMU_1361c was incubated with the *smu_137* promoter, and no mobility shifts appeared when 1361c was incubated with the *smu_136c* ORF fragment (Fig 4A). These results indicate that SMU_1361c can directly bind to the promoters of *smu_137*-*smu_141*.

**Fig 4.**
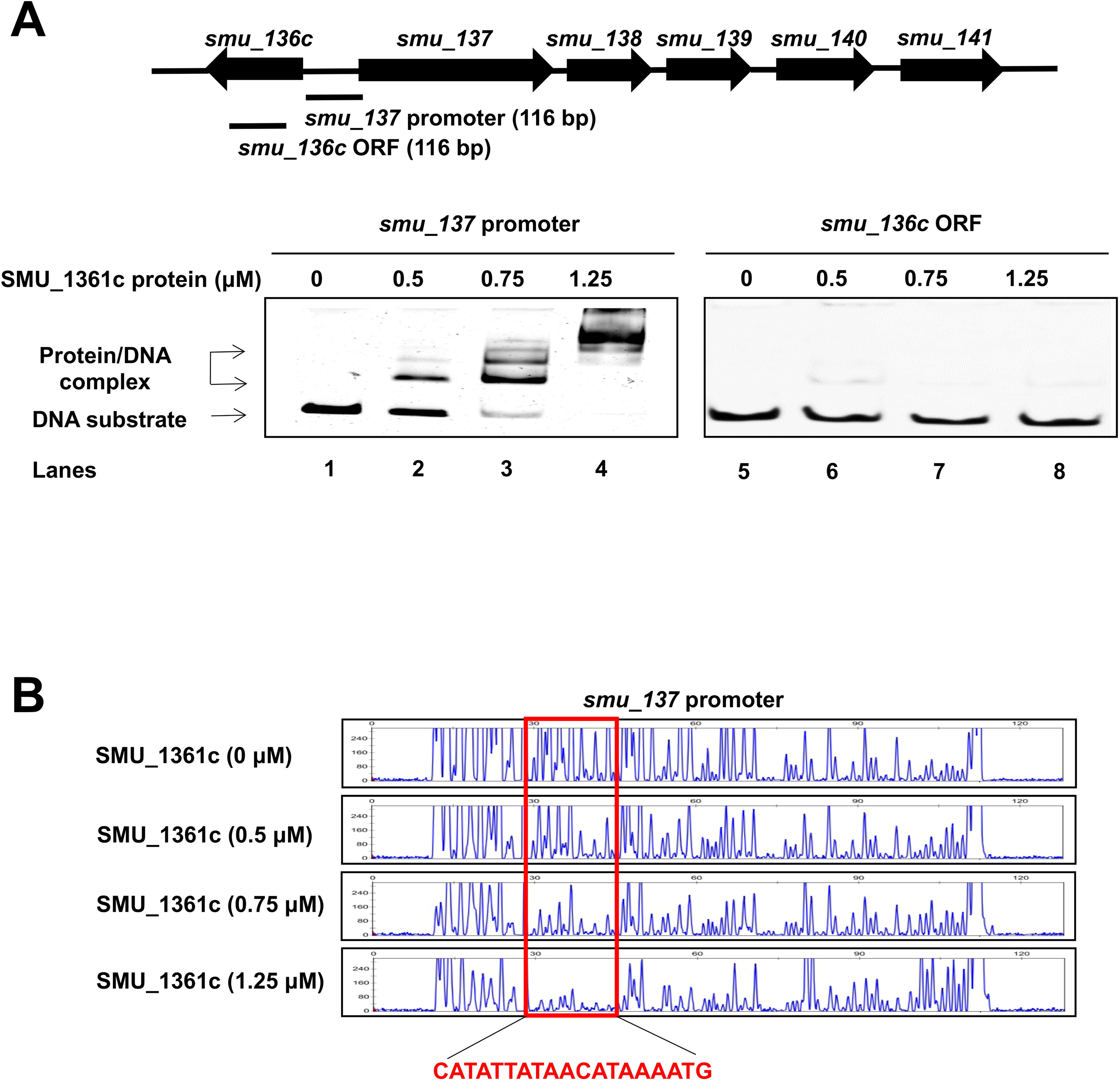
Identification of the promoter and specific DNA motif bound by SMU_1361c. (A) DNA substrates were incubated with different concentrations of SMU_1361c protein, which was analyzed using EMSA. The results show that SMU_1361c can directly bind to the promoter of *smu_137*-*smu_141* (A, lanes 1–4) and cannot bind to *smu_136c* ORF (A, lanes 5–8). (B) The *smu_137* promoter labeled with FAM was incubated with increasing concentrations of SMU_1361c protein. DNase I footprinting assay was performed to identify the specific binding motif, which was protected by SMU_1361c protein from DNase I digestion and marked in red.

### Identifying the specific binding motif of SMU_1361c

The DNase I footprinting assay was used to demonstrate the specific binding motif of SMU_1361c to the putative *smu_137* promoter region. After the *smu_137* promoter region was exposed to DNase I, the fragment containing 5′-CATATTATAACATAAAATG-3′ with high AT content was gradually protected with the increasing SMU_1361c concentrations (Fig 4B). The results showed that the specific binding motif of SMU_1361c is the palindromic sequence 5′-CATATTATAACATAAAATG-3′.

## Disscussion

Oral biofilms are microbial communities with a high density and diversity of bacterial species; these species both cooperate and compete with one another [35,36]. Many commensal bacteria, such as *S. gordonii* and *S. sanguinis*, are the initial colonizers on the tooth surface before *S. mutans* [37,38]. They create oxidative stress conditions by producing hydrogen peroxide, which gives them the ability to compete with *S. mutans* for the same ecological niche [39,40]. Therefore, the ability to tolerate oxidative stress is vital for the survival and fitness of *S. mutans* in oral biofilms. Our study results reveal that SMU_1361c, a novel TetR transcription factor, impairs the oxidative stress response by regulating the expression of multiple related virulence genes in *S. mutans* and further provide insights into the molecular mechanisms of SMU_1361c as a transcription repressor that manipulates numerous related virulence genes in *S. mutans* by directly binding to their specific promoter regions.

Since *S. mutans* lacks catalase, alternative strategies must be used to cope with the oxidative stress challenges imposed by the competing species and host [41,42]. In this study, we found that the overexpression strain UA159/pDL278-*1361c* was more sensitive to hydrogen peroxide stress than the UA159/pDL278 strain (Fig 1). In addition, the *smu_1361c* overexpression decreased the ability of *S. mutans* to compete with *S. gordonii* and *S. sanguinis* in the well-characterized interspecies interaction three-species biofilms model, signifying that the ability to tolerate oxidative stress is impaired (Fig 2). This inhibition phenotype was lost when catalase was added to the three-species biofilms, further approving that this inhibition of overexpression strain UA159/pDL278-*1361c* was a function of hydrogen peroxide generated by *S. gordonii* and *S. sanguinis*. These phenotypes were consistent with the transcriptomics analysis results of UA159/pDL278 and UA159/pDL278-*1361c*. For example, the downregulated gene clusters *smu_137*-*smu_141* is closely associated with the oxidative adaptive adaptability of *S. mutans*, which attribute to the observed defects of UA159/pDL278-*1361c* in tolerance against hydrogen peroxide (Fig 3).

The TetR family, a family of transcriptional regulators, is well represented and widely distributed amid bacteria with a DNA-binding motif, whose function is usually that of repressors [43,44]. In this study, the expression levels of the three operons associated with the oxidative stress response were downregulated when the *smu_1361c* gene was overexpressed, confirming that SMU_1361c acts as a repressor of operon expression, as seen as in other family members of *S. mutans*, such as *mubR*, *hapR*, *accR* and *frtR* [45–48]. In detail, the deletion of *mubR* and *frtR* downregulated the transcription of NRPS-PKS gene clusters and *frtP* gene, which are responsible for amplified sensitivity to oxidative stress and fluoride, respectively. *smu_1361c* may fine-tune cellular physiology during periods of oxidative stress to help preserve normal cellular homeostasis.

According to the transcriptomics analysis in this study, *smu_1361c* does not show a regulatory relationship with its neighboring genes; however, it is involved in the regulation of genes outside its operon. The EMSA results support a direct mechanism for the SMU_1361c regulation of two gene clusters *smu_137*-*smu_141.* The binding motif was identified by DNase I footprinting assays, and the results showed that a palindromic sequence with high AT content was protected by SMU_1361c, suggesting that SMU_1361c prefers binding to DNA motif with high AT content. The preference was consistent with other TetR family transcription factors, such as MubR, HapR, AccR or FrtR [45–48].

In summary, SMU_1361c, a novel TetR family transcription factor, acts as a repressor to impair the ability of *S. mutans* to survive and compete with commensal bacteria due to the sensitivity to their by-product, hydrogen peroxide. Further studies have explained that *smu_1361c* leads to significant reductions in several gene clusters involved in the oxidative stress response by binding to their specific promoter regions. Thus, an improved understanding of the mechanism by which SMU_1361c regulates the *S. mutans* oxidative stress response can strengthen our understanding to exploit interspecies interactions in the plaque biofilm for reducing dysbiosis-induced cariogenesis.

## Materials and Methods

### Bacterial strains and growth conditions

All strains, plasmids, and primers used in this study are listed in the S1 and S2 Tables. *S. mutans* UA159, *S. gordonii* DL1, and S. sanguinis SK36 were obtained from the State Key Laboratory of Oral Diseases (Sichuan University, Chengdu, China). *S. mutans* UA159 and its derivatives, *S. gordonii* DL1, and *S. sanguinis* SK36 were routinely grown in brain heart infusion (BHI) broth (Difco, Sparks, MD, United States) or on BHI agar incubated aerobically for 24 h at 37 °C. For the biofilm formation assay, 1% sucrose (wt/vol, Sigma, United States) was added to the BHI medium (BHIS). In addition, *Escherichia coli* and its derivatives were cultured in Luria-Bertani medium (BD, Sparks, MD, United States) aerobically (95% air, 5% CO_2_) with spectinomycin (1 mg/mL) or kanamycin (30 μg/mL) at 37°C, when necessary.

### Construction of overexpression strains

The genes of *smu_1361c* and *ldh* promoter regions were amplified from *S. mutans* genomic DNA by polymerase chain reaction (PCR). All the primers were designed using an online tool provided by TaKaRa (https://www.takarabio.com/learning-centers/cloning/primer-design-and-other-tools) and listed in S2 Table. The PCR products were purified and cloned into linearized *E. coli*-*Streptococcus* shuttle vector pDL278 by using an In-Fusion HD cloning kit (TaKaRa, Japan) as described previously [49]. Then, the recombined plasmid was transformed into *S. mutans* UA159 to generate overexpression strains UA159/pDL278-*1361c*, which were selected by using plates containing spectinomycin (1 mg/mL) and verified by PCR and sequencing.

### Construction of markerless in-frame deletion strains

The markerless in-frame deletion strain UA159 Δ*1361c* was made using a two-step transformation method as described previously [50]. Briefly, approximately 1 kb of the homologous sequence upstream and downstream of the *smu_1361c* open reading frame (ORF) was amplified from *S. mutans* UA159 genomic DNA using PCR with the upF/upR and dnF/dnR primers and the selection cassette IFDC2 (positive for erythromycin and negative for *p*-Cl-phe) was amplified by the *ldh*F/*erm*R primers, respectively. Then, the fragments containing the overlapping regions were ligated using the overlap extension PCR with upF/dnR primers, which were transformed into *S. mutans* UA159 and selected by using BHI agar plate containing erythromycin (12 μg/mL). For the second transformation, upstream and downstream fragments of *smu_1361c* were amplified using the primer upF/updnR and dnF/dnR and ligated using the primer upF/dnR. The ligated fragment was transformed into the first-step constructed mutant strain and selected by using plates containing *p*-Cl-phe (4 mg/mL, Sigma, United States). Finally, the markerless in-frame deletion mutant was further confirmed by PCR and sequencing.

### Quantitative reverse transcription polymerase chain reaction

Quantitative reverse transcription polymerase chain reaction (qRT-PCR) was used to quantify the expression of the *smu_1361c* gene in overexpression and deletion strains. Bacterial cells were cultured to the mid-logarithmic phase (OD_600nm_=0.5), harvested by centrifugation (4, 000 g, 4L, 10 min), and frozen in liquid nitrogen until use. To evaluate the gene expression of overexpression and deletion strains of the *smu_1361c* gene. Primer3 online tool (http://simgene.com/Primer3) was used to design the specific primer of *smu_1361c* gene. The separation, purification and reverse transcription of bacterial total RNA were carried out according to the instructions. Additionally, 16S ribosomal RNA (rRNA) gene was used as an internal control and the data were analyzed according to the 2^-ΔΔCT^ method.

### Hydrogen peroxide sensitivity assays

The hydrogen peroxide sensitivity assay was performed as follows: overnight cultures of *S. mutans* and its derivatives were washed and adjusted to the OD_600nm_ of 0.5. Four milliliters of molten BHI-soft-agar (0.6% agar, 60°C) were thoroughly mixed with 200 μL of *S. mutans* and its derivatives cell suspension and immediately poured onto the BHI agar plates, which were kept at room temperature for 30 min. Subsequently, the filter discs containing 0-, 20-, 40-, and 60-mM hydrogen peroxide were added to BHI-soft-agar plates of the test strains followed by 24 h of incubation at 37°C. Inhibition zones were measured at three different positions and averaged to a single datapoint and the ratio of inhibition zone to disc diameter was calculated.

### Bacterial composition analysis in three-species biofilms

Overnight cultures of *S. mutans* and its derivatives, *S. gordonii*, and *S. sanguinis* were harvested until OD_600nm_ to 0.5, and simultaneously inoculated (inoculum ratio of *S. mutans*/*S. gordonii/S. sanguinis*=1:1:1) in fresh BHIS or supplemented with 3 mg/mL catalase under aerobic conditions for 24 h. Then, the 24-h three-species biofilms were fixed with 4% paraformaldehyde (PFA) and labeled by species-specific fluorescent *in situ* hybridization (FISH) probes, as previously described with some modifications [33]. Biofilm images were captured using an Olympus FV3000 confocal laser scanning microscope (Olympus Corporation, Tokyo, Japan) at 60× magnification from at least five randomly selected positions of each sample. The three-dimensional structure of the three-species biofilms was reconstructed by IMARIS 7.0.0 (Bitplane, Zurich, Switzerland), and the Image-Pro Plus 6.0 (Media Cybernetics) software was used to analyze the bacterial composition.

### RNA-sequencing

For RNA-sequencing, the subcultured UA159/pDL278 and UA159/pDL278-*1361c* strains were harvested until OD_600nm_ to 0.5 under aerobic conditions, and snap frozen in liquid nitrogen until needed as described previously [51]. Total RNA from three independent samples was exacted and purified using TRIzol reagent according to the manufacturer’s instructions (Invitrogen, United States). RNA quality was determined by 2100 Bioanalyzer (Agilent, United States) and the RNA concentration was quantified using an ND-2000 instrument (NanoDrop Technologies, United States). cDNA library was constructed from enriched mRNA samples using the TruSeq RNA sample prep kit (Illumina, San Diego, CA, United States). rRNA depletion from total RNA was performed using the Ribo-Zero magnetic kit (Epicentre Biotechnologies, United States) and the mRNA was chemically fragmented to short pieces (200 nt) using a 1 × fragmentation solution (Ambion, United States) for 2.5 min at 94°C. Double-stranded cDNA was produced using the SuperScript double-stranded cDNA synthesis kit (Invitrogen) with random hexamer primers (Illumina). Then, the synthesized cDNA was subjected to end repair, phosphorylation, and “A” base addition according to Illumina’s library construction protocol. Libraries were selected for cDNA target fragments of 200 bp on 2% low-range Ultra agarose followed by PCR amplification using Phusion DNA polymerase (NEB, United States) for 15 PCR cycles. After quantification by TBS380, the paired-end RNA-seq sequencing library was sequenced with the Illumina HiSeq X Ten instrument (2 × 150-bp read length). Genes with a 2-fold difference and a P-value < 0.05 were selected as differentially expressed genes (DEGs). Functional annotation and enrichment analysis of DEGs were performed in Gene Ontology (GO) and Kyoto Encyclopedia of Genes and genomes (KEGG) databases.

### Cloning, expression, and purification of recombinant SMU_1361c protein

*smu_1361c* was amplified from *S. mutans* genomic DNA and purified, digested by *EcoR*I and *Xba*I, and cloned into the expression vector pET28a (Novagen) with an N-terminal fusion of 6 × His-tag. The reconstructed plasmid was cloned to the *Escherichia coli* strain BL21 (DE3) cells. Then, overnight cultures of the transformant were diluted at 1:20 with fresh LB medium containing 50 μg/mL kanamycin until OD_600nm_ to 0.6. After further growth with 1 mM isopropyl-β-D-thiogalactopyranoside (IPTG) at 37°C for 6 h, which induced protein expression, the cell pellets were harvested and lysed by sonification. Then, the recombinant SMU_1361c protein was purified using a His-tagged protein purification kit (Beyotime, Shanghai, China) following the manufacturer’s protocol, and concentrated by 10 kDa MWKO ultra-filtration (Millipore Amincon, Merck, Germany) from the cell debris as described previously [52]. Purified SMU_1361c concentration was determined by measuring its spectrophotometric absorbance at 280 nm, further confirmed by SDS-PAGE, and stored at -80°C for future use.

### Electrophoretic mobility shift and DNase I footprinting assays

Electrophoretic mobility shift assay (EMSA) was performed to confirm the DNA binding activity and specificity of SMU_1361c using protocols described previously with some modifications [53]. Briefly, DNA fragments containing *smu_137 promoter* (116 bp) and *smu_136c* ORF (116 bp) were amplified by PCR using the primer listed in S2 Table. Then, DNA fragments were incubated with recombinant SMU_1361c protein in the binding buffer of 50 mM Tris/HCl, 10 mM NaCl, 0.5 M Mg/acetate, 0.1 mM EDTA, and 5% (vol/vol) glycerol for 30 min on ice. DNA-protein complexes were separated by non-denaturing polyacrylamide gel electrophoresis at 110 V for 60 min. The gels were dyed with Ethidium Bromide (Thermo Scientific, Waltham, MA, US) for 20 min, and images were captured using a ChemiDoc touch imaging system (Bio-Rad, United States).

DNase I footprinting assay was used to determining the specific binding motif of SMU_1361c under similar conditions to those described for EMSA analysis. Briefly, the *smu_137* promoter region was amplified using specific primers labeled with FAM. After purification, DNA fragment (112.5 nM) was mixed with recombinant SMU_1361c (0, 2.5, 7.5, 12.5 μM), which was treated with DNase I (1 unit, Solarbio) at 37°C for 5 min. Then, the reaction was stopped using phenol chloroform, and DNA fragments were precipitated with ethanol, and eluted in 15 μL distilled water. The obtained fragment was analyzed using a 3730XL DNA analyzer (Applied Biosystems, Tsingke, China). Electropherograms were analyzed and aligned using GENEMAPPER software (Thermo Fisher Scientific, United States).

### Statistical analysis

Statistical analysis was performed with SPSS 20.0 (SPSS Software Inc, Chicago, IL, USA). Data are presented as means ± SDs from three biological replicates performed in triplicate. The student’s test was used to compare data between two groups. Analysis of variance and Tukey’s test were performed to compare data between multiple groups. *P* < 0.05 was considered statistically significant.

## Acknowledgments

We are thankful fo Jumei Zeng for their technical assistance in electrophoretic mobility shift and DNase I footprinting assays.

## Supporting information

**S1 Fig. Gene expression level of *smu_1361c* in indicated strains.** Gene expression level of *smu_1361c* was determined by qRT-PCR and caculated using the 2-ΔΔCt method with values nomalized to the reference gene 16s rRNA. The results are presented as mean ± SD (*** *P* < 0.001 or ** *P* < 0.01).

**S1 Table. Bacterial strains and plasmids used in this study.**

**S2 Table. Primers used in this study.**

## Notes

### Competing Interest Statement

The authors have declared no competing interest.

### Summary of Updates

Figure1-5 revised and Supplemental files revised.

