## Supplementary figures and images for "SMU_1361c regulates the oxidative stress response of *Streptococcus mutans*"

### S1 Fig

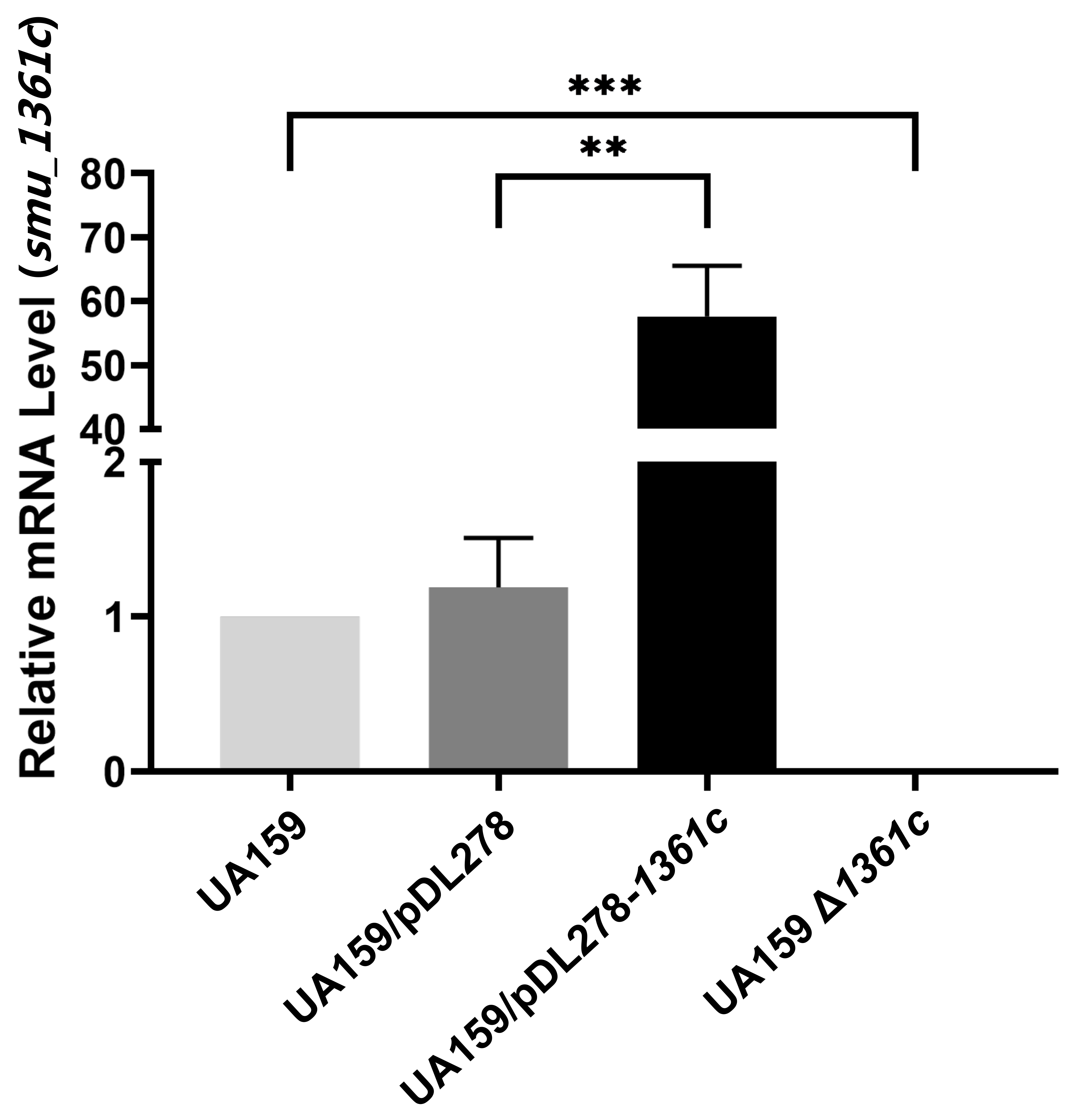
